# Minimizing the elastic energy of growing leaves by conformal mapping

**DOI:** 10.1101/2022.09.28.509914

**Authors:** Anna Dai, Martine Ben Amar

**Author notes:** Coresponding author.

## Abstract

During morphogenesis, the shape of living species results from growth, stress relaxation and remodeling. When the growth does not generate any stress, the body shape only reflects the growth density. In two dimensions, we show that stress free configurations are simply determined by the time evolution of a conformal mapping which concerns not only the boundary but also the displacement field during an arbitrary period of time inside the sample. Fresh planar leaves are good examples for our study: they have no elastic stress, almost no weight, and their shape can be easily represented by holomorphic functions. The growth factor, isotropic or anisotropic, is related to the metrics between the initial and current conformal maps. By adjusting the mathematical shape function, main characteristics such as tips (convex or concave or sharp-pointed), undulating borders and veins, can be mathematically recovered, which are in good agreement with observations. It is worth mentioning that this flexible method allows to study complex morphologies of growing leaves such as the fenestration process in Monstera deliciosa, and can also shed light on many other 2D biological patterns.

---

During morphogenesis or embryogenesis, biological species grow very slowly often creating important shape transformations at the origin of elastic stresses. In recent years, the theoretical framework of finite elasticity with multiplicative decomposition [1–3] was employed to understand and mimic these shape transformations: for instance, the development of leaves or flowers [4–6], the morphological instabilities of human organs in foetal life, including the brain cortex [7], the fingerprints [8, 9], the oesophagus mucosa [10] and the intestine villi [11–13]. For slender soft bodies with initially a rather symmetric shape, the growth may change drastically their aspect with curling and buckling [10, 14, 15], and these instabilities perfectly illustrate the successive bifurcation steps induced by the relative volume increase *G*. Thin bilayers exhibit zig-zag instabilities [12, 16–18] in the same way as fluids in Rayleigh-Bénard convection [19, 20] or localized solitonic patterns in the presence of defects [21]. However, due to the complexity of finite elasticity, most of the theoretical works describe simple highly symmetric bodies, such as thin plates or shells, with the space-independent parameter *G*, which grows slowly with time. The elastic stresses are in the order of *μV*_0_(*G −*1), *μ* being the shear modulus and *V*_0_ the initial volume: this estimate causes a significant increase of energy in the soft material if the stress relaxation and the shape remodeling are inhibited by the boundaries [3], but it may not coincide with true situations.

Recently Xiaoyi Chen *et al*. [22, 23] have proposed that the change of geometric shape is only induced by the volumetric growth *G*(*t*) without the generation of elastic stresses. This strategy implies the definition of a mapping which connects the initial position of points to their current position at time *t*, and then *G*(*t*) is determined by imposing a zero-stress condition. Their derivation is rather technical and their examples are based on initial circular geometry. Nevertheless, they derive a variety of stress free mathematical geometries that mimic five biological patterns.

With the same objective but limiting ourselves to two-dimensional thin samples of arbitrary initial and final shapes, like fresh leaves, we propose a general formalism based on conformal mapping techniques. Indeed, regular planar close curves *∂*Ω limiting a domain Ω can be related to the unit circle by an holomorphic function according to the Riemann theorem. This function also associates the points inside Ω to the Riemann disc, and it gives a way to construct the geometric deformation field during the growth process. Then the shape will evolve from cartesian coordinates to a rectangular curvilinear system of coordinates. D’Arcy Thompson [24] made the hypothesis that these coordinates represent the velocity of shaping. Although conformal mapping is not explicitly mentioned in the original work, the pictures drawn in Ref.[24] concerning the growth and evolution between neighbor species strongly suggest such hypothesis. The idea behind conformal mapping for growing leaves was revisited more recently [25] and also tested experimentally. For example, K. Alim et al. [26] predict the local displacement field of petunia and tobacco leaves through a conformal mapping, which is rather consistent with their experimental results.

The aim of this work is to prove that conformal mappings can recover the shape of leaves without generating elastic stress and can determine the growth laws, independently of the nonlinear elasticity model. Herein, different shapes of common leaves are selected, with a special focus on the Monstera deliciosa family.

## The formalism

We consider a 2D formalism where the leaf thickness remains constant during the growth without deformation in the thickness direction. Growth is a very slow process so that the deformations adjust immediately to the growth. The initial leaf shape is represented by Ω_0_ and the material points by **X**, the current shape at time *t* becomes Ω_*t*_ with the current point coordinates **x**. The geometric deformation gradient is the second order tensor defined by: 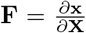, and **F** = **F**_**e**_**G** where **G** is the growth and **F**_**e**_ the elastic tensor [1]. The right Cauchy tensor **C** only depends on **F**_**e**_ and is given by 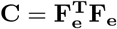. Using the curvilinear coordinates of the initial configuration with *Z* = *X* + *iY* = *F*_1_(Ξ) = *F*_1_(*μ*+*iη*), we choose *z* for the current configuration, such as *z* = *x*+*iy* = *F*_2_(*k*(*μ*)+*il*(*η*)) where conformal mapping is preserved only if *k*(*μ*) = *μ* and *l*(*η*) = *η. F*_2_(*μ* + *iη*) is determined by the outer leaf shape *∂*Ω_*t*_ which corresponds to *μ* = *μ*_0_ = *k*(*μ*_0_) at the time *t* of observation. Introducing *k*(*μ*) and *l*(*η*) for *z* simply broadens the ensemble of mappings between the two domain boundaries *F*_1_ and *F*_2_.

Local growth rate of living tissues can be inhomogeneous (dependent on the coordinates *μ* and *η*) and anisotropic, which explains the tensorial mathematical representation of **G**. To respect the leaf geometry, this tensor **G** must be diagonal: **G** = *diag* (*g*(*μ, η*)*/p*(*μ, η*), *p*(*μ, η*)*g*(*μ, η*)), where *p*(*μ, η*) is the growth anisotropy coefficient, and *g*(*μ, η*)^2^ is the local volumetric growth at time *t*. For simplicity, we suppress the *μ, η* and *t* dependence in p and g functions. Then the elastic tensor becomes:

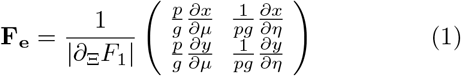

For incompressible living species, the third invariant *I*_3_ = det **F**_*e*_ = 1, so imposing the local growth eigenvalue *g*:

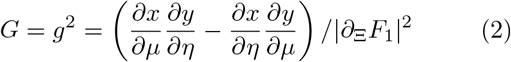

A stress free configuration requires the cancellation of the first invariant *I*_1_: *I*_1_ = Trace **C** *−*2 = 0 which, with the constraint *I*_3_ = 1, leads to:

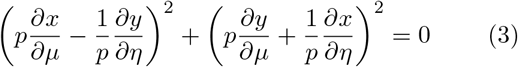

If *p* = 1, the growth is isotropic, and Eq.(3) recovers the Cauchy relations implying that the current configuration *z* is an holomorphic function. If *p* ≠ 1,the growth is anisotropic and *p*^2^ = *l*^*′*^(*η*)*/k*^*′*^(*μ*). The derivation can be found in the Supplemental Material (SM), Sec.I. In finite elasticity, the elastic strain depends on invariants (invariant under a rotation) traditionally called *I*_1_, *I*_2_ and *I*_3_ [27, 28], also eventually on pseudo-invariants *I*_4_ and *I*_5_ for transversely isotropic fibrous materials [29]. All these invariants are functions of the right Cauchy tensor **C** which is reduced to the unity tensor. So as long as Eq.(3) is satisfied, the growth process will generate no stress, even in the case of anisotropic growth.

As a conclusion, for 2D materials, it exists an infinite space of stress-free conformal maps associated to a growth tensor satisfying simultaneously Eq.(2) and Eq.(3). Incompressibility is not mandatory since material compressibility is controlled by *I*_3_ that is equal to unity in our case. In the following, we focus on fresh leaves which are extremely diverse in nature with quasi-planar shapes independently of the connection to the branch. The vein size depends on the species, most of them have a central prominent and rigid vein with a network of weaker lateral veins [30]. Having its own characteristics, each species requires an adaptation of our model to recover its shapes, which is perfectly doable with holomorphic functions. In particular, when very rigid veins appear, a partition following the big veins may be necessary and the modeling should be applied piece by piece. Hereafter, we focus on botanic traits, such as the tip, the margin, and the existence of internal holes. The knowledge of the initial and current shape contours specifies both functions *F*_1_ and *F*_2_ and also the growth density *g*. However, despite the mathematical proofs of existence of such functions, their precise determination remains a challenge in practice requiring to solve a rather difficult inverse problem [31, 32], and the accuracy of numerical methods depends strongly on the complexity of the domain geometry. Therefore, our choice will consist in summing a restricted number of hyperbolic cosine functions −*i*Σ_*k*_*b*_*k*_(*t*)*S*^*k*^ with *S* = cosh [*a*(*μ* + i*η*)], the coefficient *b*_*k*_ being obtained by simple fitting of the contour. In practice, 3 modes *k* were sufficient to mimic a variety of leaf shapes during their growth. The schematic diagram is shown in Fig.1.

**FIG. 1.**
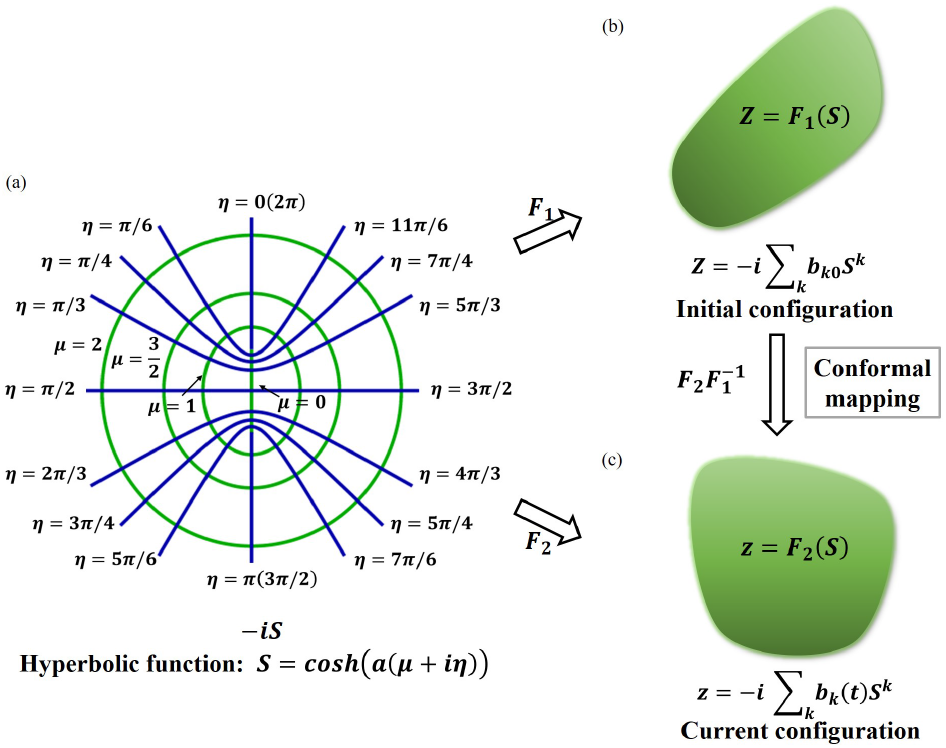
Schematic diagram: (a) *μ* and *η* are the curvilinear coordinates, *μ* = *μ*_0_ corresponds to a green curve and *η* = *η*_0_ to a blue curve. (b) Parameters *b*_*k*0_ depend on the leaf contour at *t* = 0, (*F*_1_), (c) parameters *b*_*k*_(*t*) are determined by the contour at time *t*, (*F*2).

## Tips and margins of leaves

Tips play a major role in physical growth processes in various fields such as dendritic growth [33], viscous fingering [34], fractures [35–37] and filamentary organisms [38]. Tips are often considered as being responsible for not only the growth dynamics but also the stability of the global shape. However, leaves [39] exhibit almost all kinds of shapes at the tip such as sharp-pointed, convex or concave, see Fig.(2)(a)-(c), the last case being much less common. Also, leaf margins are rather diverse being either smooth such as lily leaves or undulated such as apple leaves. All these morphologies can be recovered with our formalism based on the expansion in powers of *S*, by fixing the central vein at *μ* = 0 and the outer contour at *μ* = *μ*_0_. The coefficient *a* characterizes the plant species and the parameters *b*_*k*_ at initial and final time *t* are adjusted to the observed contours, then generating both functions *F*_1_ and *F*_2_ (see SM, Sec. SII). Nevertheless, tip with a central dip or undulating margins require an additive correction *S*_*c*_ = *−ic*(*t*)*Se*^*d*(*t*)(*μ*+*iη−*0.6)^ to our expansion. The period and the amplitude of oscillations are easily controlled by *c*(*t*) and *d*(*t*) respectively, see Fig.3, where we focus on the margins of the Jujube and the White Sapote leaves. For the Robinia shown in Fig.2(f), the term *S*_*c*_ plays two roles, i.e., controlling tip and margin. Once the shape of the leaves is obtained with enough accuracy, the volumetric growth can be evaluated using Eq.(2). In particular, in Fig.2(d)-(f), color variation from dark green (weak growth amount) to light green or yellow indicates the heterogeneity of the growth intensity, less pronounced for the most rounded leaf (Fig.2(e)).

**FIG. 2.**
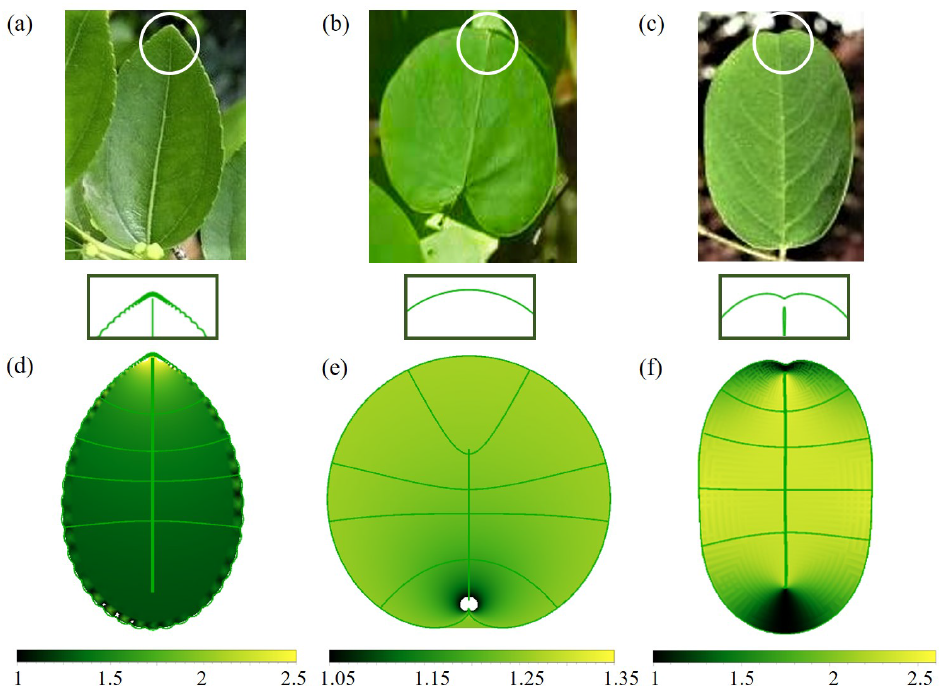
Natural leaves: (a) Jujube leaf with sharp tip and oscillatory border. (b) Redbud leaf with rounded tip and smooth border. (c) Robinia pseudoacacia leaf with concave tip and smooth border. Mathematical images: (d)-(f) simulated by shape function (see SM, Sec. SII). Level of green colors shows the local growth density in different regions, lighter color indicating more growth intensity. Jujube leaf (a) and (d) grows more at the tip while the top and bottom of Robinia pseudoacacia leaf (c) and (f) have a weak level of growth. Redbud leaf (b) and (e) shows a relatively uniform growth in the entire area, except near the petiole.

**FIG. 3.**
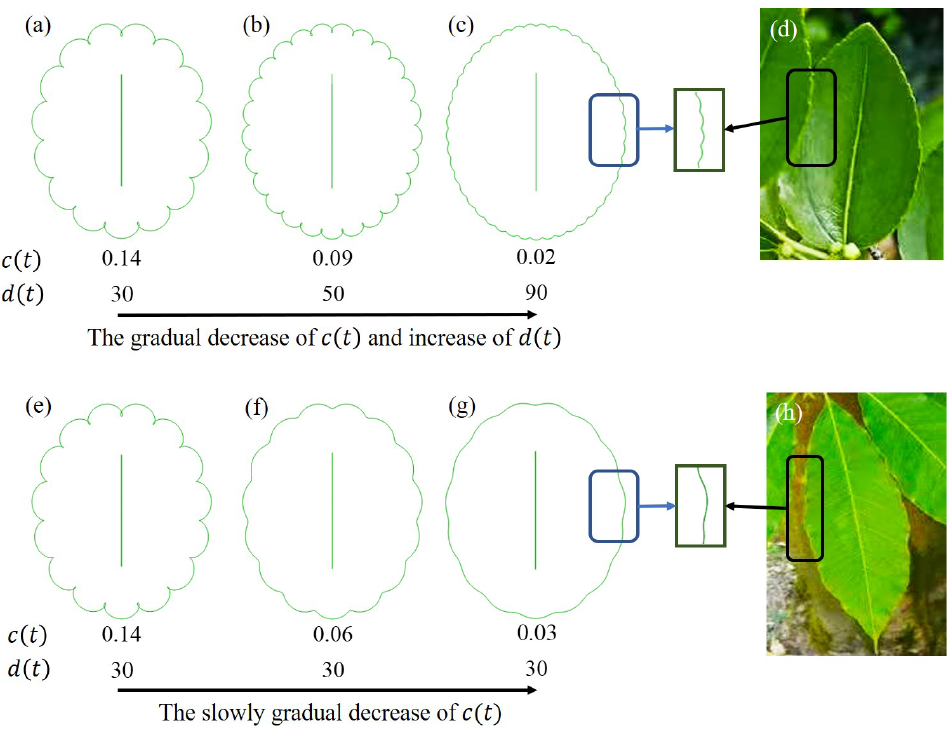
Different leaf borders, shape function of all contours: *z* = *−ib*_1_*S* + *S*_*c*_, where *μ*_0_ = 0.6, *a* = 2 and *b*_1_ = 2. (a)-(c): Boundary shapes changing with decreasing values of *α*(*t*) and *β*(*t*). (d) Jujube leaves showing similar edges with (c). (e)-(g) Boundary shapes evolving with only decreasing values of *α*(*t*). (h) White Sapote leaves show similar edges as (g).

## Blade perforation

Surprisingly, the initiation and evolution of holes in leaves also called fenestration is a rather rare event across the plant world, and it mostly happens in the family of Monstera deliciosa. It has become a very fashionable decorative vine nowadays. The blade perforation remains difficult to interpret in terms of adaptive function to its natural environment, that consists in tropical forests, and different hypotheses have been considered such as water uptake or sun flecks [40]. There are some biological evidences that the blade perforation is generated by a regulated program of cell death [41] and in this case, we must take the hole distribution as a matter of fact occurring in rather big leaves (more than 10 cm): physics and mechanics cannot explain or justify their existence and distribution. Indeed, even in the same vine, the leaves have no symmetry: some present holes on both sides of the central veins, some only on one side, and the number of holes per leaf is highly variable. Most of the time, the perforation is not visible and happens when the leaf is still inside the sheath of an old one, see SM. The emergence of a hole in a mature leaf is a rare event but not impossible, in this case the central zone of the domain between two lateral veins becomes thinner and thinner in the middle, and ultimately one hole appears. The typical time scale for this event is about the month.

We have noticed the similarity of hole shapes with viscous fingers or bubbles in Hele-Shaw cell [42]. From the viewpoint of modeling, they have in common to be generated with conformal fields, of course due to totally different physical reasons. The studies more connected to our work concern series of steady bubbles of velocity *U* travelling periodically in an infinite linear Hele-Shaw cell. When surface tension is neglected, their shapes are defined by 2 (for symmetric and centered bubbles, [43]) or 7 parameters (for non symmetric bubbles,[44–46]), each of them allowing a time-dependent adjustment. The periodic flow field is Laplacian, satisfies the imposed boundary conditions on the two sides of a rectangle which corresponds to one period of the flow and is limited by the parallel horizontal walls of the experimental cell. Such rectangle can be easily mapped to a domain enclosed by two lateral veins (defined by *η*), the central vein and the outer contour. At the bubble boundary, the pressure vanishes which is also relevant to the elastic fenestration problem without stress. Calling *ζ* the Riemann unit disc coordinates, *z*_*b*_ the position of an arbitrary point in the flow, Burgess and Tanveer establish the following relation [43]:

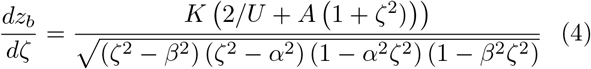

where *K* and *A* are constants determined by the three parameters *U* and *α* and *β*, they are responsible for the length and width of the hole through Eq.(4). The transformation process and the mapping functions are displayed in Fig.4(b). The leaf shape functions are fully determined at two different times of their evolution and are shown in Fig.4(a), in which the maximum growth rate *G* is predicted near the petiole while the tip corresponds to a minimal growth. These solutions have symmetric holes (for-aft, up-down) and centered in the middle, but in reality, holes may appear closer to the edge where they broaden, see Fig.4(c). In this case, they lose the for-aft symmetry and display finger shapes, the previous approach [43] ceases to be valid and more complex shape functions involving 8 parameters are required [44–46], see SM. The mapping process between the bubble and the hole is the same for symmetric or asymmetric case, see Fig.4(b) and Fig.4(d). In addition, a peculiar variety of Monstera exhibits a finger facing a small hole which reminds viscous fingering experiments in Hele-Shaw cell [46–48]. Fig.4(e) displays an example of such leaves, comparing to our mathematical shapes at two different times. To sum up, our formalism applies to any kind of planar leaves. The shape complexity may require elaborated mapping functions that can be found either in the literature or in classical specialized books [49–51]. Numerical methods have also been established [31, 32, 52]. The amount of growth, isotropic or anisotropic, is obtained as displayed in Fig.4 and in SM.

**FIG. 4.**
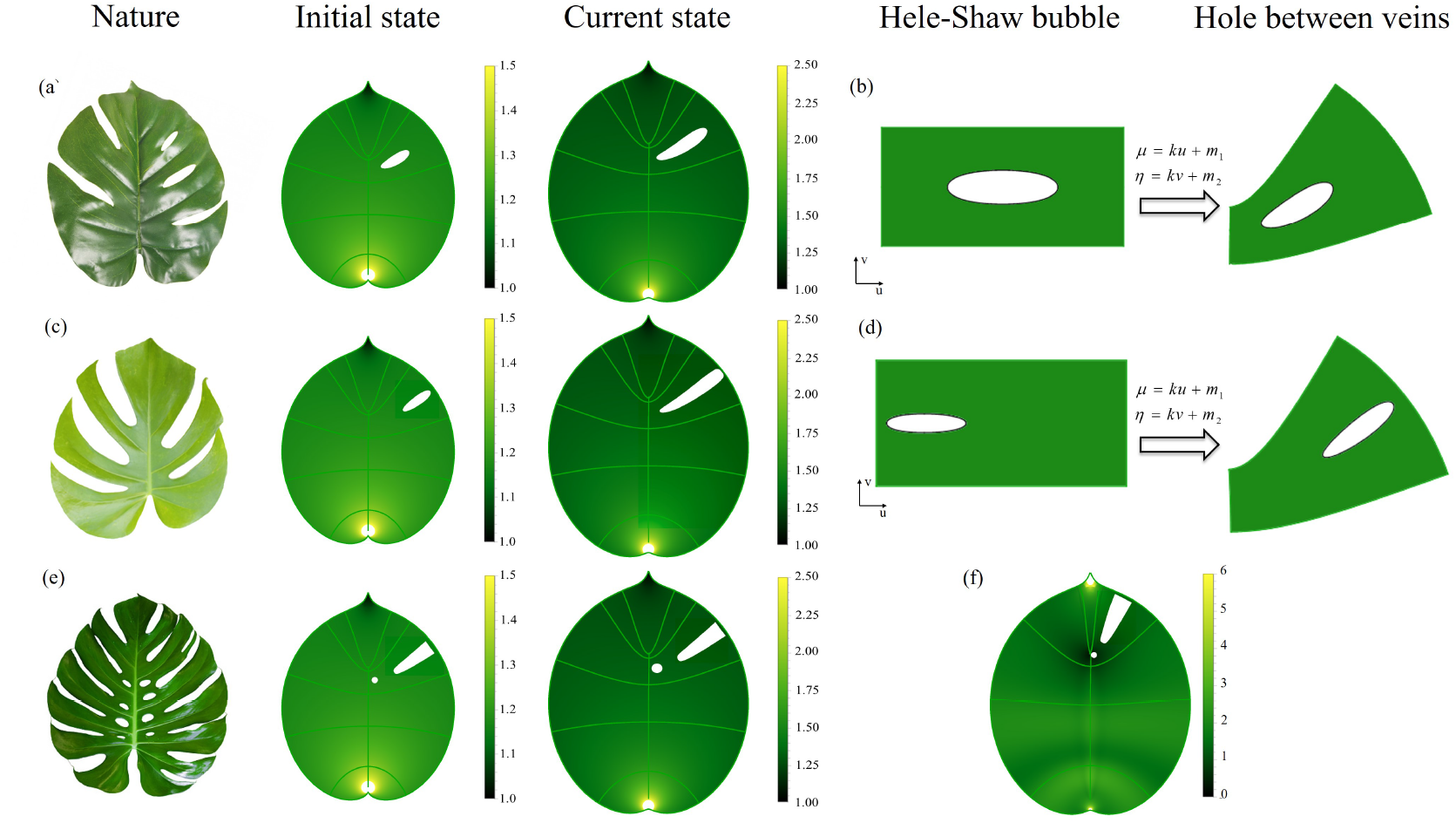
(a) On left, natural Monstera deliciosa leaf with holes located either at the center (on top right) or connected to the border. On the same level on top, initial and current state of a mathematical leaf with one hole, located at the center in a chosen area. (b) A rectangle including a hole on left mapped into a specific curvilinear rectangle with boundaries defined by *μ* and *η*. The selected conformal mapping is a solution of the periodic Darcy flow with periodic and symmetric bubbles. Selection of constant *k* and *c*_2_ are made in relation with (a). (c) Natural Monstera leaf with ‘fingers’ like shape holes. The initial and current state of mathematical leaf with asymmetrical hole. (d) Conformal mapping generating an asymmetric bubble. The hole is closer to the outer boundary. (e) Natural Mexico Monstera leaf with holes and fingers and the initial and current state of mathematical leaf with two holes. (f) Anisotropic growth of the mathematical leaves with two holes, the functions *k*(*μ*) and *l*(*η*) are detailed in [49], notice the changes of vein position between (e) and (f).

## Leaf vein

Veins provide structure and support to leaves while also playing a vital role in transporting water and nutrients to the leaf blade. The region closer to the major vein may have access to more nutrients, which will be evenly transmitted into lateral veins with a nutrient content equivalent in each location [30]. Such unequal nutrient distribution is responsible for an anisotropic growth process represented in the footnote [53]. It further leads to a change of the leaf vein locations and consequently of the hole shape, as shown by the comparison of Fig.4 (f) and (e). Besides in the anisotropic case, the growth rate *G* is much different from the isotropic one and turns out to be more homogeneous in the whole leaf, except near the petiole and the tip.

## Conclusion

Among all mappings possible for the shape evolution of a 2D elastic sample, we demonstrate that conformal or quasi-conformal mappings have the advantage to eliminate the elastic stresses independently of the elastic material properties. Contrary to other cases studied recently [3, 5, 6], leaves without exterior loading and growing in a quiet environment sustain this approach. In this letter, we exploit the hypothesis of conformal mapping [24–26] on plant leaves, recovering the boundary and further obtaining the displacement field which establishes the growth kinematics. Our method extracts information not only on the cell proliferation which is often restricted to the nutrient penetration but also on the biological complexity, such as tissue remodeling. [40]. This formalism allows to evaluate the growth accumulation in case of isotropy or anisotropy. Veins can also be simulated and their relationship with nutrient contents can be established. Understanding the stress-free morphological evolution induced by growth is not limited to the morphogenesis of leaves or other biological tissues, but can also shed light on the design of new biomimetic soft devices.

## Supporting information

Supplement of main manuscript

## I. ACKNOWLEDGEMENTS

The authors acknowledge the support of ANR (Agence Nationale de la Recherche) under the contract MecaTiss (ANR-17-CE30-0007) and the contract EpiMorph (ANR-2018-CE13-0008). Anna Dai acknowledges the support of the CSC (China Scholarship Council), file No. 201906250173.

## Notes

### Competing Interest Statement

The authors have declared no competing interest.

