## Supplement of main manuscript for "Minimizing the elastic energy of growing leaves by conformal mapping"



(Dated: September 28, 2022)


### I. STRESS-FREE GROWTH

We assume that the initial shape of the leaf is given by:

$$Z = F_1(\Xi) = F_1(\mu + \mathrm{i}\eta) \quad \text{so} \quad \Xi = \mu + i\eta \quad \text{and} \quad G_1 = F_1^{-1} \tag{1}$$

where $i^2 = -1$, $F_1$ represents the mapping between cartesian coordinates and curvilinear coordinates of the interior of the leaf, $\mu$ and $\eta$ are conformal coordinates and $\mu = \mu_0$ gives the initial outer shape. Here, we restrict to leaf shapes simply described by a enough regular contour. We will explain later domains with holes which must be treated with appropriate methods [1–3]. Knowing the outer boundary, it is always possible to define the function $F_1$ from the boundary, which is not always an easy task and this mapping is unique, for regular contour. To take the inverse of an holomorphic function $F_1$ is always doable formally and gives $G_1$ also an holomorphic function. As a consequence, we get the following Cauchy relations:

$$\frac{\partial \mu}{\partial X} = \frac{\partial \eta}{\partial Y} \quad \text{and} \quad \frac{\partial \mu}{\partial Y} = -\frac{\partial \eta}{\partial X} \tag{2}$$

In morpho-elasticity, growth is represented by a tensor $\mathbf{G}$ which is inhomogeneous most of the time (that means space dependent) and also anisotropic. We assume that the growth respects the leaf geometry so the tensor $\mathbf{G}$ is diagonal and reads:

$$\mathbf{G} = \begin{bmatrix} \frac{1}{p(\mu,\eta)} g(\mu, \eta) & 0 \\ 0 & p(\mu, \eta)\, g(\mu, \eta) \end{bmatrix} \tag{3}$$

where $p(\mu, \eta)$ is the growth anisotropy coefficient and $\mathrm{Det}(\mathbf{G}) = g(\mu, \eta)^2$ is the volumetric growth at the time of observation. When the leaf grows, we assume that the displacement of each point is given by the following mapping $F_2$ such that:

$$z = F_2(k(\mu) + il(\eta)) \tag{4}$$

where $z$ represents the new coordinates $(x, y)$. This mapping is not conformal but it gives for $x$ and $y$ the relationship with the initial shape configuration such as:

$$\begin{cases} x = \frac{1}{2}\left[F_2(k(\mu) + il(\eta)) + \overline{F_2}(k(\mu) - il(\eta))\right] \\ y = \frac{1}{2i}\left[F_2(k(\mu) + il(\eta)) - \overline{F_2}(k(\mu) - il(\eta))\right] \end{cases} \tag{5}$$

According to the main hypothesis of the morpho-elasticity theory, the geometric gradient of deformation $\mathbf{F}$ and the elastic gradient of deformation tensor $\mathbf{F_e}$ are related by : $\mathbf{F} = \mathbf{F_e G}$ [4]. Written in these conformal coordinates, these tensors become:

$$\mathbf{F} = \frac{1}{|\partial_\Xi F_1|} \begin{bmatrix} \frac{\partial x}{\partial \mu} & \frac{\partial x}{\partial \eta} \\ \frac{\partial y}{\partial \mu} & \frac{\partial y}{\partial \eta} \end{bmatrix} \quad \text{and} \quad \mathbf{F_e} = \frac{1}{|\partial_\Xi F_1|} \begin{bmatrix} \frac{p}{g}\frac{\partial x}{\partial \mu} & \frac{1}{pg}\frac{\partial x}{\partial \eta} \\ \frac{p}{g}\frac{\partial y}{\partial \mu} & \frac{1}{pg}\frac{\partial y}{\partial \eta} \end{bmatrix} \tag{6}$$

The first physical constraint concerns the incompressibility of the sample which imposes:

$$\mathrm{Det}(\mathbf{F_e}) = 1 \Longleftrightarrow \frac{\partial x}{\partial \mu}\frac{\partial y}{\partial \eta} - \frac{\partial x}{\partial \eta}\frac{\partial y}{\partial \mu} = g^2 |\partial_\Xi F_1|^2 \tag{7}$$

In addition, a stress-free configuration imposes $I_1 = Tr\left(\mathbf{F_e}^{\mathbf{T}}\mathbf{F_e}\right) - 2 = 0$ and we derive:

$$p^2\left(\frac{\partial x}{\partial \mu}\right)^2 + p^2\left(\frac{\partial y}{\partial \mu}\right)^2 + \frac{1}{p^2}\left(\frac{\partial x}{\partial \eta}\right)^2 + \frac{1}{p^2}\left(\frac{\partial y}{\partial \eta}\right)^2 = 2g^2|\partial_\Xi F_1|^2 = 2\left(\frac{\partial x}{\partial \mu}\frac{\partial y}{\partial \eta} - \frac{\partial x}{\partial \eta}\frac{\partial y}{\partial \mu}\right) \tag{8}$$

A simple reorganization of Eq.(8) leads to:

$$\left(p\frac{\partial x}{\partial \mu} - \frac{1}{p}\frac{\partial y}{\partial \eta}\right)^2 + \left(p\frac{\partial y}{\partial \mu} + \frac{1}{p}\frac{\partial x}{\partial \eta}\right)^2 = 0 \tag{9}$$

so we get:

$$p\frac{\partial x}{\partial \mu}-\frac{1}{p}\frac{\partial y}{\partial \eta}=0 \quad \text{and} \quad p\frac{\partial y}{\partial \mu}+\frac{1}{p}\frac{\partial x}{\partial \eta}=0 \tag{10}$$

If $p=1$ (isotropic growth), Eq.(10) recovers the Cauchy relations and implies that $F_2$ is an holomorphic function. If $p\neq 1$, an anisotropic growth process may also generate a stress free configuration. The coefficient of anisotropy for a description ruled by $F_2$ (see Eq.(5)) is obtained with:

$$\frac{\partial x}{\partial \mu}=k'(\mu)\,\frac{{F_2}'+\overline{F_2}'}{2},\quad \frac{\partial x}{\partial \eta}=l'(\eta)\,i\frac{{F_2}'-\overline{F_2}'}{2},\quad \frac{\partial y}{\partial \mu}=k'(\mu)\,\frac{{F_2}'-\overline{F_2}'}{2i},\quad \frac{\partial y}{\partial \eta}=l'(\eta)\,i\frac{{F_2}'+\overline{F_2}'}{2} \tag{11}$$

For simplicity, we do not detail the variable of each function and the symbol prime ($'$) means the first derivative with respect to the natural variable as defined in Eq.(5). So introducing the values of the partial derivatives given by Eq.(11) into Eq.(10) gives:

$$p^2=\frac{l'(\eta)}{k'(\mu)} \tag{12}$$

### II. COMPLEX REPRESENTATIONS OF LEAVES

In the main text, four different types of leaves are simulated, showing different properties. Knowing their outer border, their respective shape functions are derived from the hyperbolic function $S=\cosh\left(a(\mu+i\eta)\right)$, eventually with a correction $S_c$. We consider 2 different times. The first one called the initial time or time t=0 corresponds to the situation where the leaf is yet planar and its shape characteristics rather well defined, the second time concerns the situation where obviously the surface of the leaf has increased and perhaps some new features concerning eventually the tip or the margin appear. Due to the growth, the coefficients between the two configurations will evolve in time and will reveal the observed shapes of mature leaves at time $t$. In the cases considered in the main manuscript, we have firstly found the contour function of the Jujube leaf, which is similar to an oval, with sharp tips and jagged edges, and it reads:

$$z=-i\left(b_1(t)S+b_2(t)S^2+b_3(t)S^3+c\,(t)\,e^{d(t)(\mu+i\eta-0.6)}S\right) \tag{13}$$

Redbud leaf resembles an inverted heart shape, with rounded tip and smooth border:

$$z=-i\left(b_1(t)S+b_2(t)S^2\right) \tag{14}$$

Robinia pseudoacacia leaf has a concave tip and smooth border suggesting:

$$z=-i\left(b_1(t)S+b_2\,(t)\,S^2+c\,(t)\,e^{d(t)(\mu+i\eta-0.6)}S\right) \tag{15}$$

Monstera leaf is the same as Redbud leaf, their petiole is concave, but the overall leaf is slightly longer, narrower and the tip is more pronouced:

$$z=-i\left(b_1(t)S-b_2\,(t)\,S^2+0.1(\mathrm{Log}\,(S-S_0))^{1.09}\right) \tag{16}$$

where $S_0=\cosh\left(a\left(\mu_v+I\eta_v\right)\right)$, $(\mu_v,\eta_v)$ being located at the vertex coordinate of the function $z=-i\left(b_1(t)S-b_2\,(t)\,S^2\right)$. $a$ is a constant, and $b_k\,(t)$, $c(t)$ and $d(t)$ are functions of $t$. Except for the Monstera, the data are obtained by simple guess in Table S$I$. For the Monstera leaf, the data between two time intervals of $14$ days are derived by fit of the outer contour at time $t=0$, $t_1$ (leave 1) and $t_2$ (leave 2). In the following, we restrict first on the Redbud case, then on the Monstera case.

### III. DETERMINATION OF THE GROWTH CHARACTERISTICS

In this section, we detail the method to reach the growth parameters as $g(\mu,\eta)$ and $p(\mu,\eta)$ for the Redbud leaf. From Eq.(14), we choose the initial representation as :

$$\begin{aligned} X&=-b_{10}\sin(a\eta)\sinh(a\mu)-2b_{20}\cos(a\eta)\cosh(a\mu)\sin(a\eta)\sinh(a\mu)\\ Y&=b_{10}\cos(a\eta)\cosh(a\mu)+b_{20}\{\cos(a\eta)\cosh(a\mu)\}^2-b_{20}\{\sin(a\eta)\sinh(a\mu)\}^2 \end{aligned} \tag{17}$$

| Leaf Name | $\mu_0$ | a | $b_{10}$ | $b_1(t)$ | $b_{20}$ | $b_2(t)$ | $b_{30}$ | $b_3(t)$ | $c_0$ | $c(t)$ | $d_0$ | $d(t)$ |
|---|---|---|---|---|---|---|---|---|---|---|---|---|
| Jujube Leaf | 0.51 | 1 | 0.8 | 1 | 0.55 | 0.6 | 0.16 | 0.16 | 2.42 | 6.58 | 60 | 70 |
| Redbud Leaf | 1 | 1.1 | -0.62 | -0.7 | -0.5 | -0.22 | - | - | - | - | - | - |
| Robinia pseudoacacia leaf | 0.7 | 1 | 1.5 | 1.75 | 0.28 | 0.3 | - | - | 0.11 | 0.09 | 3 | 3 |
| Monstera leaf 1 | 0.8 | 1.2 | 1.7 | 1.8 | 0.58 | 0.6 | - | - | - | - | - | - |
| Monstera leaf 2 | 0.8 | 1.2 | 1.8 | 1.9 | 0.6 | 0.63 | - | - | - | - | - | - |

TABLE.S I. Parameters of leaf functions at initial time and at time $t$. For Monstera, two times of observation are listed.independently of $t = 0$.

and the current representation as:

$$\begin{aligned} x &= -b_1(t)\sin(a\eta)\sinh(a\mu) - 2b_2(t)\cos(a\eta)\cosh(a\mu)\sin(a\eta)\sinh(a\mu) \\ y &= b_1(t)\cos(a\eta)\cosh(a\mu) + b_2(t)\{\cos(a\eta)\cosh(a\mu)\}^2 - b_2(t)\{\sin(a\eta)\sinh(a\mu)\}^2 \end{aligned} \tag{18}$$

The volumetric growth density parameter $g(\mu,\eta)^2$, for isotropic growth has been established in Eq.(2) of the main text, and is displayed in Fig.(2) and Fig.(4) of the main text, with green colors of variable intensity according to the values of $g^2$. Once we fix the initial configuration at time $t = 0$ with the same representation (which is not mandatory), we get the following volumetric growth rate:

$$g^2 = \frac{b_1(t)^2 + 2b_2(t)^2\cos(2a\eta) + 4b_1(t)b_2(t)\cos(a\eta)\cosh(a\mu) + 2b_2(t)^2\cosh(2a\mu)}{{b_{10}}^2 + 2b_{20}^2\cos(2a\eta) + 4b_{10}b_{20}\cos\cos(a\eta)\cosh(a\mu) + 2{b_{20}}^2\cosh(2a\mu)} \tag{19}$$

The same method can be applied for finding the growth rate which determines the stress-free pattern for the other leaves. All the parameters of leaf functions are shown in Table SI. Fig.(2) of the main text shows three kinds of leaves: Jujube, Redbud and Robinia pseudoacacia leaf. In Fig.(4) (a)(c)(e) devoted to Monstera, the initial state leaf uses the parameters of the Monstera leaf 1 to calculate the growth. Current state leaf and Fig.(4) (f) use the parameters of the Monstera leaf 2.

To illustrate the method for anisotropic growth, we define $k(\mu) = \mu$ and $l(\eta) = (2\pi)^{-2}e^{\frac{(\eta^2-1)}{(2\pi)^2}} + \eta - 0.1$ according to Eq.(3) of the main manuscript, defining the current configuration:

$$z = ib_1(t)\cosh\left[a\left(\mu + il\left(\eta\right)\right)\right] + ib_2(t)\cosh^2\left[a\left(\mu + il\left(\eta\right)\right)\right] \tag{20}$$

According to Eq.(12), the anisotropic coefficient $p$ is given by $p^2 = 1 + (8\pi^4)^{-1}\eta e^{\frac{\eta^2}{(2\pi)^2}-1}$, and the volumetric growth rate becomes:

$$g^2 = \frac{l'(\eta)\{\cos(2al(\eta)) - \cosh(2a\mu)\}\left\{b_1(t)^2 + 2b_2(t)^2(\cos(2al(\eta)) + \cosh(2a\mu)) + 4b_1(t)b_2(t)\cos(al(\eta))\cosh(a\mu)\right\}}{\{\cos(2a\eta) - \cosh(2a\mu)\}\{b_{10}^2 + 2b_{20}^2\cos(2a\eta) + 4b_{10}\mathrm{b}_{20}\cos(a\eta)\cosh(a\mu) + 2b_{20}^2\cosh(2a\mu)\}} \tag{21}$$

where the parameters are the same as the ones for the isotropic growth. The mathematical leaves are shown in Fig.(S1) and we can observe that the position of the leaf veins has changed with the growth anisotropy. So growth anisotropy gives a way to modify the position of the lateral veins if their positions change between the two times of observation.

### IV. HELE-SHAW BUBBLES VERSUS HOLES IN MONSTERA

The Monstera leaf fenestration requires specific tools for its representation. Our strategy consists in isolating the area where the hole is located maintaining the original whole shape given by Eq.(16). We look for a conformal mapping which involves the domain between the central axis, two consecutive veins and the outer frontier of the leaf defined by $\mu_0 = 0.8$. The strategy is obvious if a complex mapping is known for a complex potential function defined in a stripe with one or two holes. The potential flow representing a bubble or a finger travelling in a linear Hele-Shaw cell can be a good candidate. Indeed, the hydrodynamic flow being governed by the Darcy law with incompressibility leads to a Laplacian pressure field, the shape bubble appearing as a line of zero-pressure. The classical work of a unique finger [5] or bubble [6] in the linear infinite channel geometry has been solved by Saffman and Taylor a long time ago, and the more recent extension to a periodic set of bubbles has been discovered later, giving a way to solve a major difficulty: the domain of interest in our case has a finite length and we must link this domain to a unique period of the stationary flow.

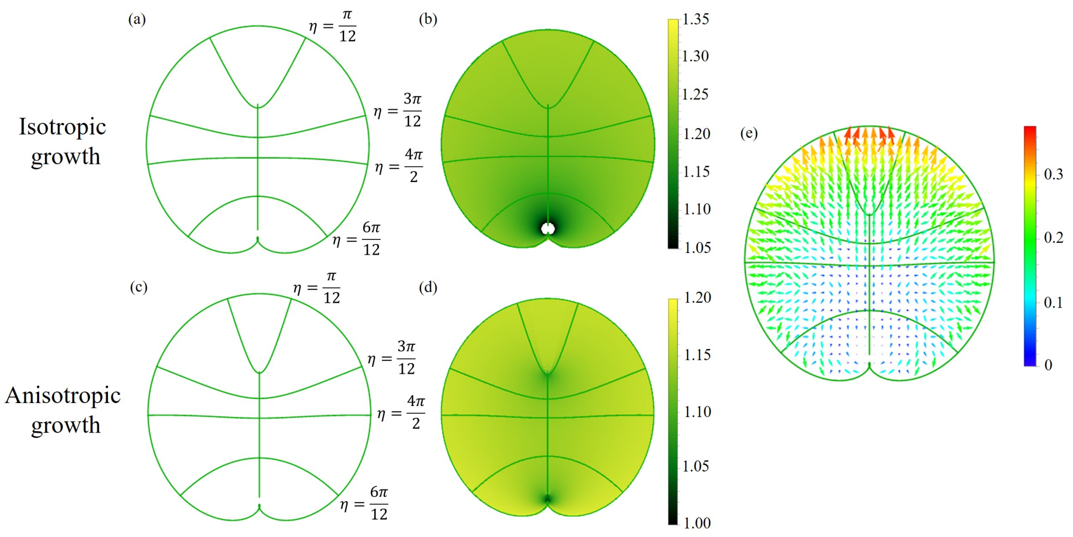


Fig.S 1. (a) Current outer contour corresponding to $\mu_0 = 1$ of the isotropic growth. Veins of constant value $\eta = \eta_0$. (b)Volumetric isotropic growth coefficient in green colors. Intensity varies between 1.05 and 1.35. (c) Current outer contour corresponding to $\mu_0 = 1$ of the anisotropic growth. Veins of constant value $\eta = \eta_0$. For anisotropic growth $p^2 = 1 + (8\pi^4)^{-1}\eta e^{\frac{\eta^2}{(2\pi)^2}-1}$. Comparing to the (a), the position of veins changed. (d) Volumetric anisotropic growth coefficient in green colors. Intensity varies between 1 and 1.2. (e) Displacement of the velocity field visualized with arrows.

#### A. Symmetric hole

We first focus on the symmetric hole/bubble located in the middle of the domain. D. Burgess and S. Tanveer [7] have derived a three-parameter family of exact solutions defined in the unit half-disc, see Fig.(S2)(a)-(b). We select this conformal mapping $z_b(\zeta)$ determined by:

$$\frac{dz}{d\zeta} = KF(\zeta)\left(2/U + A + A\zeta^2\right) \quad \text{with} \quad F(\zeta) = \frac{1}{\sqrt{(\zeta^2-\beta^2)(\zeta^2-\alpha^2)(1-\alpha^2\zeta^2)(1-\beta^2\zeta^2)}} \tag{22}$$

where $z_b(\zeta) = u + iv$, and $(u, v)$ are the coordinates in the physical plane of the Hele-shaw cell moving with a bubble with the velocity $U$. $\alpha$, $\beta$ and $U$ are the 3 degrees of freedom, while the constants $K$ and $A$ are determined via $\alpha$, $\beta$ and $U$:

$$f_n(\alpha,\beta) = \int_\alpha^\beta t^n F(t)dt; n = 0, 2, \quad K = \frac{U-1}{f_0 - f_2}, \quad \text{and} \quad A = \frac{1}{f_0 + f_2}\left(\frac{f_0 - f_2}{U-1} - \frac{2f_0}{U}\right) \tag{23}$$

These parameters are within the following ranges: $1 < U < \infty$, $0 < \alpha < \beta < 1$, $K > 0$ and $-1/U < A < \infty$.

As shown in Fig.(S2)(a) and (b) similar to the Fig.(2) in the Ref.[7], six points from $A$ to $E$ represent the one-to-one correspondence points before and after the mapping. We divide them into four parts, line segment CD, EF, DE and half bubble AB, and then we use Eq.[22] for piecewise integration to obtain the Fig.(S2)(b), which shows a half-bubble in the rectangle. It is worth noting that in the integration we need to specify the length $L$ and width of the rectangle. The width is constant in this mapping and equal to 1, L is determined by:

$$g_n(\alpha,\beta) = \int_{-\alpha}^{\alpha} t^2 F(t)dt; n = 0, 2 \quad \text{and} \quad K[(2/U + A)g_0 + Ag_2] = L \tag{24}$$

By means of the symmetry in Fig.(S2)(b), a complete bubble is obtained, and an additive conformal mapping is required for the transformation of the bubble into a hole located between two lateral curved veins. By letting $\alpha = 0.79$, $\beta = 0.808$, $U = 3$ for Fig.(S2)(c) and $\alpha = 0.9$, $\beta = 0.935$, $U = 3.5$ for Fig.(S2)(d), the original hole shape at two different times (green figures) are obtained. Then the transformation can be achieved by a simple linear relationship, $\mu = ku + m_1$ and $\eta = kv + m_2$, where constants $k = \pi/16$, $m_1 = 0$ and $m_2 = \pi/16$, two cases are obtained with holes in different areas, and their details are shown in Fig.(S2)(c)-(d).

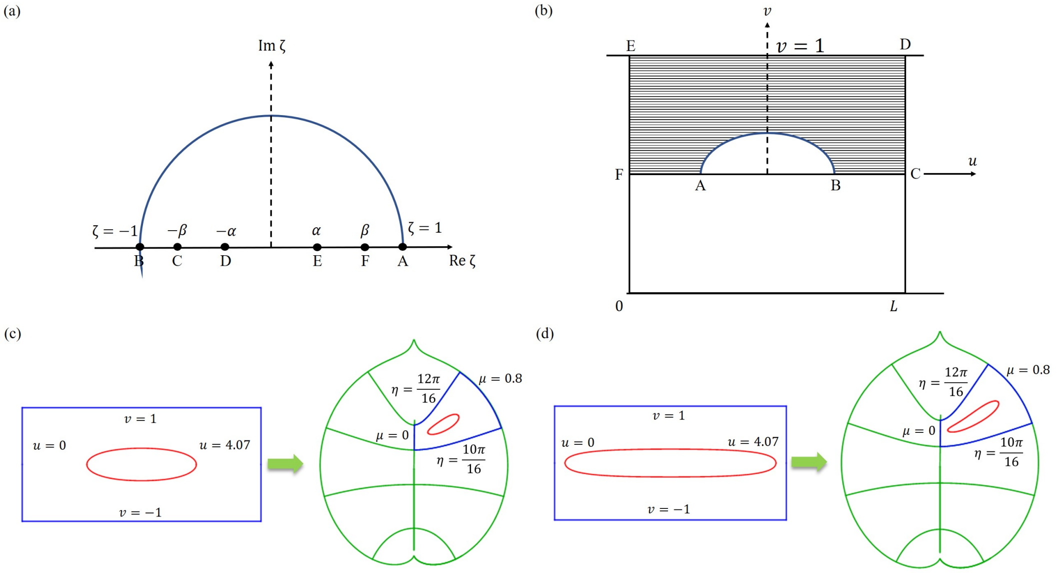


Fig.S 2. In (a) the flow region in the $\zeta$ plane, in (b) the physical $z_b$ plane from [7]
. In (c) and (d) original and transformed holes at time $t = 0$ and time $t$ in a growing leaf.

#### B. Asymmetrical hole/bubble

Noticing that, in the Monstera deliciosa, the shape of holes looks like more a viscous finger issued from the outer boundary, we consider now a process where the left-right symmetry is not preserved. Such solutions and extension of the previous case [7] have been explicited in different works by G. Vasconcelos and his collaborators [8, 9] and D. Crowdy [10, 11], and following the formulation developed in Ref. [12], see also Fig.(S3)(a)-(b), the function $z_b(\zeta)$ reads:

$$z_b\left(\zeta\right) = -\frac{1}{U}\left[W(\zeta) - \widetilde{W}(\zeta)\right] \tag{25}$$

where $W(\zeta)$ and $\widetilde{W}(\zeta)$ are defined by :

$$W(\zeta) = \mathrm{K}\int_{\zeta_0}^{\zeta} \frac{P\left(q^2\frac{\zeta}{\beta_1}, q\right) P\left(q^2\frac{\zeta}{\beta_3}, q\right)}{\sqrt{\prod_{k=1}^4 P\left(\frac{\zeta}{\alpha_k}, q\right)}} d\zeta \quad \text{and} \quad \widetilde{W}(\zeta) = \widetilde{\mathrm{K}}\int_{\alpha_4}^{\zeta} \frac{P\left(q^2\frac{\zeta}{\beta_2}, q\right) P\left(q^2\frac{\zeta}{\beta_4}, q\right)}{\sqrt{\prod_{k=1}^4 P\left(\frac{\zeta}{\alpha_k}, q\right)}} d\zeta \tag{26}$$

the constants K and $\widetilde{\mathrm{K}}$ being given by:

$$\mathrm{K}^{-1} = \frac{1}{w\left(U-V\right)}\int_{\theta_3}^{\theta_4} \frac{P\left(q^2\frac{e^{i\theta}}{\beta_1}, q\right) P\left(q^2\frac{e^{i\theta}}{\beta_3}, q\right)}{\sqrt{\prod_{k=1}^4 P\left(e^{i(\theta-\theta_k)}, q\right)}} e^{i\theta} d\theta \quad \text{and} \quad \widetilde{\mathrm{K}}^{-1} = -\frac{1}{wV}\int_{\theta_3}^{\theta_4} \frac{P\left(q^2\frac{e^{i\theta}}{\beta_2}, q\right) P\left(q^2\frac{e^{i\theta}}{\beta_4}, q\right)}{\sqrt{\prod_{k=1}^4 P\left(e^{i(\theta-\theta_k)}, q\right)}} e^{i\theta} d\theta \tag{27}$$

The angle $\theta_i$ satisfies the following sequence such that $0 < \theta_1 < \theta_2 < \theta_3 < \theta_4 \leq 2\pi$, $\theta_k = \arg\left(\alpha_k\right)$, see Fig. (S3)(a). The function $P\left(\zeta, q\right)$ is related to the first Jacobi Theta function $\Theta_1\left(\zeta, q\right)$:

$$P\left(\zeta, q\right) = -\frac{ie^{-\tau/2}}{Cq^{1/4}}\Theta_1\left(i\tau/2, q\right) \tag{28}$$

where $\tau = -\log\left(\zeta\right)$, and $C$ is a positive constant which will not appear in the following. $\alpha$ and $\beta$ must satisfy some relationships as:

$$\alpha_1\alpha_2\alpha_3\alpha_4 = 1 \tag{29}$$

$$\beta_1\beta_3 = q^2, \quad \beta_3 = \overline{\beta_1} \tag{30}$$

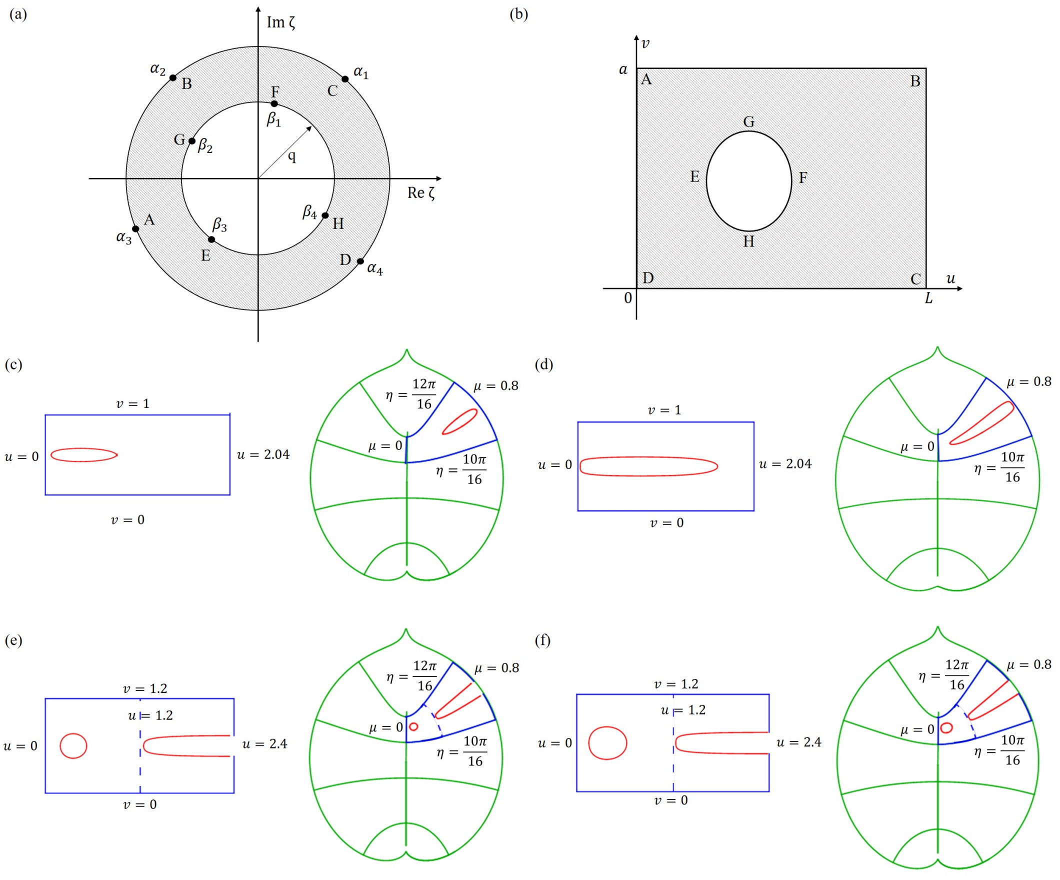


Fig.S 3. In (a) the flow region in the $\zeta$ plane, in (b) $z_b$ plane. In (c) and (d), original and transformed hole at time $t = 0$ and $t$ in the $z_b$ plane. (e)(f) Original and transformed holes at $t = 0$ and $t$ in a growing leaf.

$$\beta_2\beta_4 = -q^2, \quad \beta_4 = -\overline{\beta_2} \tag{31}$$

The imposed up and down symmetry of the bubbles (but not left and right symmetry) causes additional relations on $\alpha_i$:

$$\begin{aligned} \alpha_2 &= -\overline{\alpha_1} \\ \alpha_4 &= -\overline{\alpha_3} \end{aligned} \tag{32}$$

Then, for such solution, there are eight parameters to determine the final position and the bubble shape, the velocity $U$, $V$ $\theta_1$, $\theta_3$, $\beta_1$, $\beta_2$, $q$ and $a$ in the calculation. For simplicity, we always define $\theta_3 = \pi$, $\beta_1 = \pi/2$, $V = 1$, so $\theta_4 = 0\,(2\pi)$ and $\beta_3 = 3\pi/2\,(-\pi/2)$. Now, the bubble is obtained by five parameters $\theta_1$, $\beta_2$, $q$ and $a$, $U$. These parameters are within the following range: $0 < \theta_1 < \pi/2$, $0 < \beta_2 < \pi$, $0 < q < 1$ and $0 < w < +\infty$, $1 < U < +\infty$.

As before, a conformal mapping is required for the transformation of the bubble into a hole located between two lateral veins. By assuming $\theta_1 = 1.547$, $\beta_2 = 0.35$, $q = 0.4$, $w = 1$, $U = 4.5$ and $\theta_1 = 1.41$, $\beta_2 = 0.505$, $q = 0.68$, $w = 1$, $U = 4.5$, the original hole shape in two different states are obtained (see Fig.(S3)(c) and (d)). The transformation can be also achieved by the simple linear relationship, $\mu = -u\pi/(8a) + m_1$ and $\eta = -v\pi/(8a) + m_2$. For the process in Fig.(S3)(c), the constants are $m_1 = 4/5$ and $m_2 = 3/4$, and for the process in Fig.(S3)(d), $m_1 = 4/5$ and $m_2 = 3\pi/4$.

For the two hole examples, we still use the asymmetric bubble function to obtain a small bubble close to the left border and then get a finger-shaped bubble near the right border. The image of the finger-shaped bubble is shifted to the left by the length L, which is the length of the small bubble cell. We treat it as a whole and map it between two lateral veins. For the initial state as shown in Fig.(S3)(e), we define $\theta_1 = 1.24$, $\beta_2 = 0.26$, $q = 0.3$, $w = 1.2$, $U = 2$ to get the small bubble, and $\theta_1 = 0.82$, $\beta_2 = 1.5$, $q = 0.6$, $w = 1.2$, $U = 4.5$ to get the finger-shaped bubble. Current state is shown in Fig.(S3)(f), we define $\theta_1 = 1.22$, $\beta_2 = 0.32$, $q = 0.4$, $w = 1.2$, $U = 2$ to get the small bubble, and $\theta_1 = 0.674$, $\beta_2 = 1.3$, $q = 0.66$, $w = 1.2$, $U = 4.5$ to get the finger-shaped bubble. Linear transformation is the same as before, $\mu = -u\pi/(8w) + m_1$ and $\eta = -v\pi/(8w) + m_2$, the constants being $m_1 = 0$ and $m_2 = 5\pi/8$ for both processes.

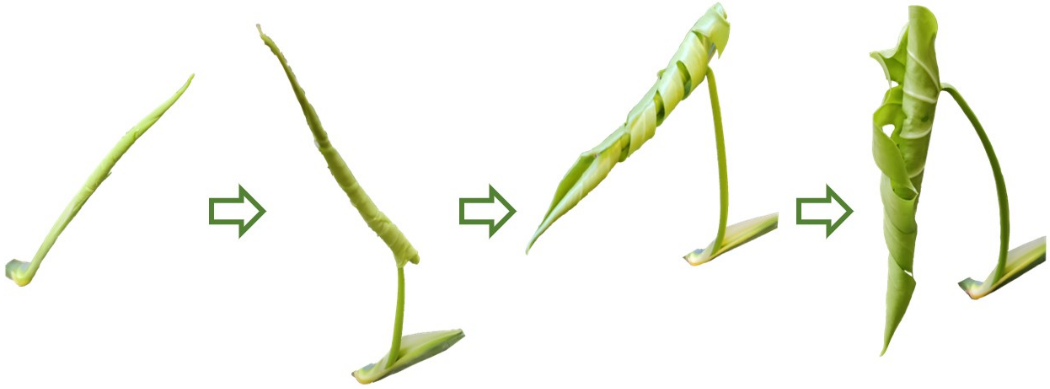

Fig.S 4. Monstera leaf in early stages.

### V. IMAGES OF YOUNG MONSTERA

In the main text, we mentioned the young Monstera leaves are inside a sheath, and they will expand over time, as shown in Fig.(S4). It takes about one month for the young leaf to get planar (which may be our initial time), but after, growth will continue during years.
